# Mechanisms of *Igf2* inhibition in thymic epithelial cells infected by coxsackievirus CV-B4

**DOI:** 10.1101/2020.01.10.902684

**Authors:** Hélène Michaux, Aymen Halouani, Charlotte Trussart, Chantal Renard, Hela Jaïdane, Henri Martens, Didier Hober, Vincent Geenen

## Abstract

Epidemiological studies have evidenced a link between type 1 diabetes (T1D) and infections by enteroviruses, especially with coxsackievirus B4 (CV-B4). CV-B4 is able to infect human and murine thymic epithelial cells (TECs) and, in a murine TEC line, we have shown that the diabetogenic strain CV-B4 E2 decreases transcription of insulin-like growth factor 2 gene (*Igf2*), coding for the self-peptide of the insulin family. Here we show that in CV-B4 infection of mice alters *Igf2* transcripts isoforms in TECs, followed by a decrease of pro-IGF2 precursor in the thymus. CV-B4 infection of a murine TEC line decreases *Igf2* P3 promoter activity by targeting the region −68 to −22 upstream of the transcription start site (TSS) whereas *Igf2* transcripts stability is not affected, pointing towards a regulation of *Igf2* transcription. Our data also show that CV-B4 decreases IL-6/STAT3 signaling *in vitro*. This study provides new knowledge about the regulation of intrathymic *Igf2* transcription by CV-B4 and reinforces the hypothesis that CV-B4 infection of the thymus could break central self-tolerance of the insulin family by decreasing *Igf2* transcription and IGF2 presentation in thymus epithelium.

**IMPORTANCE:** Coxsackievirus B4 represents one of the most important environmental factors associated to type 1 diabetes, autoimmune disease for which no curative treatment exist. The diabetogenic strain Coxsackievirus B4 E2 was previously shown to decrease *Igf2* expression, important player for central tolerance towards insulin, in a thymic epithelial cell line. The understanding of *Igf2* regulation mechanisms during coxsackievirus B4 infection represents an interest for the understanding of central tolerance development but also for *Igf2* transcriptional regulation itself, still poorly understood.

Here we demonstrate that, some transcripts isoforms of *Igf2* are also decreased in thymic epithelial cells *in vivo*. Moreover, we show that this decrease is induced by an alteration of specific regions of *Igf2* P3 promoter and may be linked by a decrease of STAT3 signaling. *In fine* we hope that this work could lead to future therapies leading to reprogramming central tolerance towards β cells antigens via *Igf2* expression.

## INTRODUCTION

The presentation of neuroendocrine self-peptides by thymic epithelial cells (TECs) plays an essential role in programming immune self-tolerance to neuroendocrine functions and a defect in this process is the earliest event in the pathogenesis of autoimmune diseases such as type 1 diabetes (T1D) (1). Insulin-like growth factor 2 (IGF2) is the self-peptide of the insulin family and *Igf2* transcription is absent in the thymus of an animal model of T1D, the diabetes-prone Bio-Breeding (DP-BB) rat (2). Furthermore, *Igf2* expression is required for the establishment of complete immune self-tolerance of insulin (3).

Enteroviruses and especially coxsackievirus B (CV-B) are among the most important environmental factors that have been linked to T1D (4). *Enterovirus* genus is part of the *Picornaviridae* family. Enteroviruses are non-enveloped small viruses, composed of a single-strand positive RNA in an icosahedric capsid and are transmitted by orofecal route. The so-called diabetogenic strain CV-B4 E2 has been isolated from a child died from ketoacidosis after an acute T1D onset (5). CV-B4, frequently detected in T1D patients, has a tropism for pancreatic insulin-secreting β cells in Langerhans islets and various mechanisms were proposed to explain the induction of autoimmune T1D by enteroviruses (4).

For several years we have been investigating whether CV-B4 could infect the thymus and disturb its crucial function in the programming of central self-tolerance to the insulin family and to islet β cells. Following oral and intra-peritoneal inoculation, CV-B4 also infects the thymus, which leads to abnormal T-cells differentiation (6–9) and this was also observed in murine and human thymic fetal organ cultures (10, 11). Also, CV-B4 is able to induce a persistent infection of primary cultures of human TECs and to modulate their profile of cytokine secretion (12). CV-B4 persistently infects the MTE4-14 cell line, a TEC line derived from neonatal mice (13), and this induces a drastic decrease in *Igf2* transcription and IGF2 production while *Igf1* transcription was much less affected (14).

In this study, we investigated the effects of CV-B4 E2 infection on *Igf2* transcription using an enrichment method of TECs (15, 16) from an outbred susceptible mice strain (9). We also explored mechanisms of the regulation of *Igf2* transcription in MTE4-14 cells induced by a short CV-B4 infection.

## RESULTS

### CV-B4 E2 decreases *Igf2* transcripts and pro-IGF2 expression in TECs *in vivo*

*Igf2* is mainly found in TECs (CD45-), which encompass only a few percent of the thymic population. We decided to use for each thymus a depletion of thymocytes based on their positive CD45 expression to enrich in CD45^−^ TECs (Fig. 1A and 1B). TECs (CD45^−^EpCAM^+^) were enriched in average 62-fold in CD45^−^ sorted cells and accordingly, *total Igf2* mRNA relative level was in average 318-fold higher in CD45^−^ sorted cells (TECs) compared to total thymic cells (Fig. 1C).

**Fig. 1.**
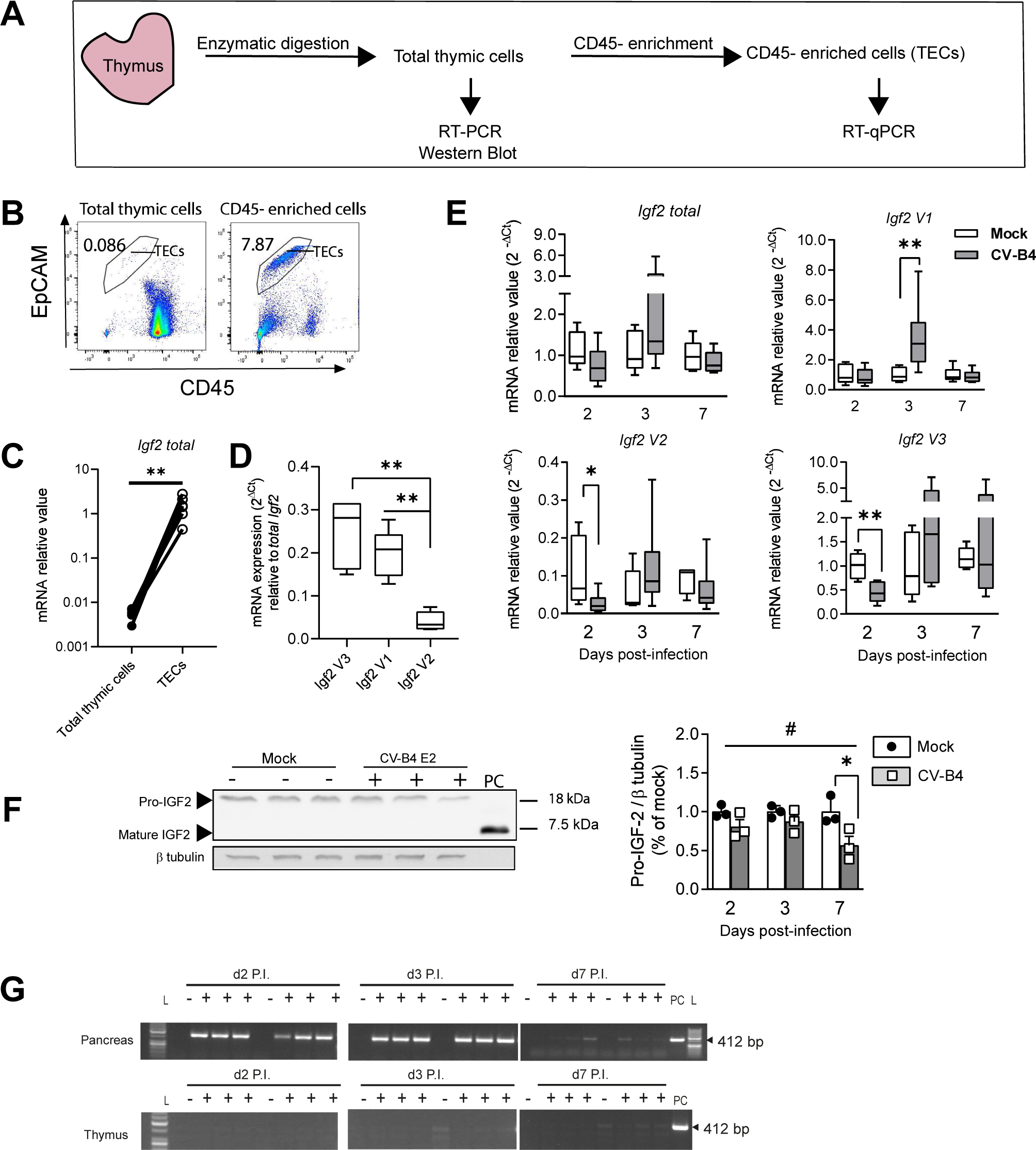
Effect of CV-B4 E2 on murine thymic *Igf2* mRNA isoforms and its precursors *in vivo.* **(A)** Schematic thymus processing for isolation of total thymic cells (unsorted cells) and CD45^−^ enriched cells (TECs). **(B)** Flow cytometry analysis of CD45 enriched TECs (EpCAM^+^CD45^−^) *versus* unsorted total thymic population. Numbers indicate the percentage of TECs population **(C)** Relative mRNA expression of *total Igf2* mRNA in unsorted total thymic population and in matched enriched TECs fraction (CD45-). Mock uninfected mice at day 2 P.I. were used; *n* = 6. **(D)** Relative mRNA expression of *Igf2 V3, V1 and V2* mRNA transcripts isoforms in CD45^−^ enriched TECs in mock uninfected mice at day 3 P.I. For each sample, relative mRNA expression of *Igf2* transcripts isoforms was normalized to the corresponding *total Igf2* mRNA expression; *n =* 6 **(E)** Relative mRNA expression of *Igf2* mRNA isoforms in mock and in CV-B4 E2 infected mice; box-and-whisker plots extend from minimum to maximum values, with lines at medians; *n* = 5-6 **(F)** Left panel, representative Western blot of IGF2 and its precursors at day 7 P.I. in mock (−) and in CV-B4 E2 infected mice (+). Independent biological samples are represented. PC, purified mature IGF2. Right panel, relative quantification of pro-IGF2 in mock and in CV-B4 E2 infected mice; histogram represents mean of relative value ± SD; *n* = 3. **(G)** Representative agarose gel electrophoresis of one-step RT-PCR products of CV-B4 E2 genome in digested thymus (total thymic cells) and in matched pancreas, in CV-B4 E2 (+) or in mock uninfected mice (−). Independent biological samples are shown. L, ladder; PC, MTE4-14 cells infected with CV-B4 E2. **(C)** Student’s paired *t* test, \*\**p* < 0.01 **(E-F)** Student’s *t* test, \*\**p* < 0.01 and \**p* < 0.05; **(D; F)** one-way ANOVA, \*\**p* < 0.01; #*p* < 0.05.

We investigated *Igf2* expression by RT-qPCR in TECs. It is known that murine *Igf2* gene contains three main promoters (P1, P2, P3) which give rise to three transcript isoforms differing only in their first exon (5’UTR): *Igf2 V3* (*Igf2* P3), *Igf2 V1* (*Igf2* P2), and *Igf2 V2* (*Igf2* P1) (17). *Igf2* Variant 3 *(Igf2 V3)* and *Igf2* Variant 2 mRNA *(Igf2 V2)* isoforms, main and minor *Igf2* transcripts isoforms respectively (Fig. 1D), were significantly decreased at 2 days post-infection (P.I.) in infected mice. However, *Igf2* Variant 1 *(Igf2 V1)* was on the opposite upregulated by CV-B4 E2 after 3 days P.I., and *total Igf2* in TECs was not significantly decreased in CV-B4 E2 inoculated mice. At 7 days P.I., relative expression for each isoform returns to comparable level to mock-inoculated mice (Fig. 1E).

Next, IGF2 protein level and its precursors were analyzed in total thymic cell population (containing both thymocytes and TECs) by Western Blot. Mainly pro-IGF2 precursor form (18 kDa, dominant isoform) was detected whereas no mature IGF2 was observed. Interestingly, compared to mock-inoculated mice, pro-IGF2 level was significantly decreased during the course of the infection, especially at day 7 P.I (Fig. 1F). Collectively, these results suggest that CV-B4 is able to decrease *Igf2* at mRNA and protein level in the thymus. Of note, despite evident sign of CV-B4 infection detected in all pancreases sampled from CV-B4 inoculated mice, CV-B4 viral RNA was surprisingly not detected within thymic cells (Fig. 1G).

### CV-B4 E2 decreases *Igf2* transcripts and pro-IGF2 expression in MTE4-14 cell line

Among *Igf2* transcripts, only *Igf2 V1* and *Igf2 V3* mRNA are detected by RT-PCR in MTE4-14 cell line (Fig. 2A) with a large dominant expression of *Igf2 V3* (Fig. 2B). During the course of CV-B4 _MOI = 0.05_ infection, RT-qPCR results show a significant and gradual decrease of *total Igf2*, starting at day 2 P.I. Moreover, among *Igf2* mRNA transcripts, both *Igf2 V3* and *Igf2 V1* were decreased of 74% at day 3 P.I. in CV-B4 _MOI = 0.05_ infected cells (Fig. 3A). Regulator of cell cycle *Tp53,* and apoptosis regulators *Birc5* and *Bax,* previously shown to be modulated during enterovirus or by others single-stranded positive RNA viruses (18–20), were not altered during the time course of infection with CV-B4 _MOI = 0.05_ (Fig. 3B).

**Fig. 2.**
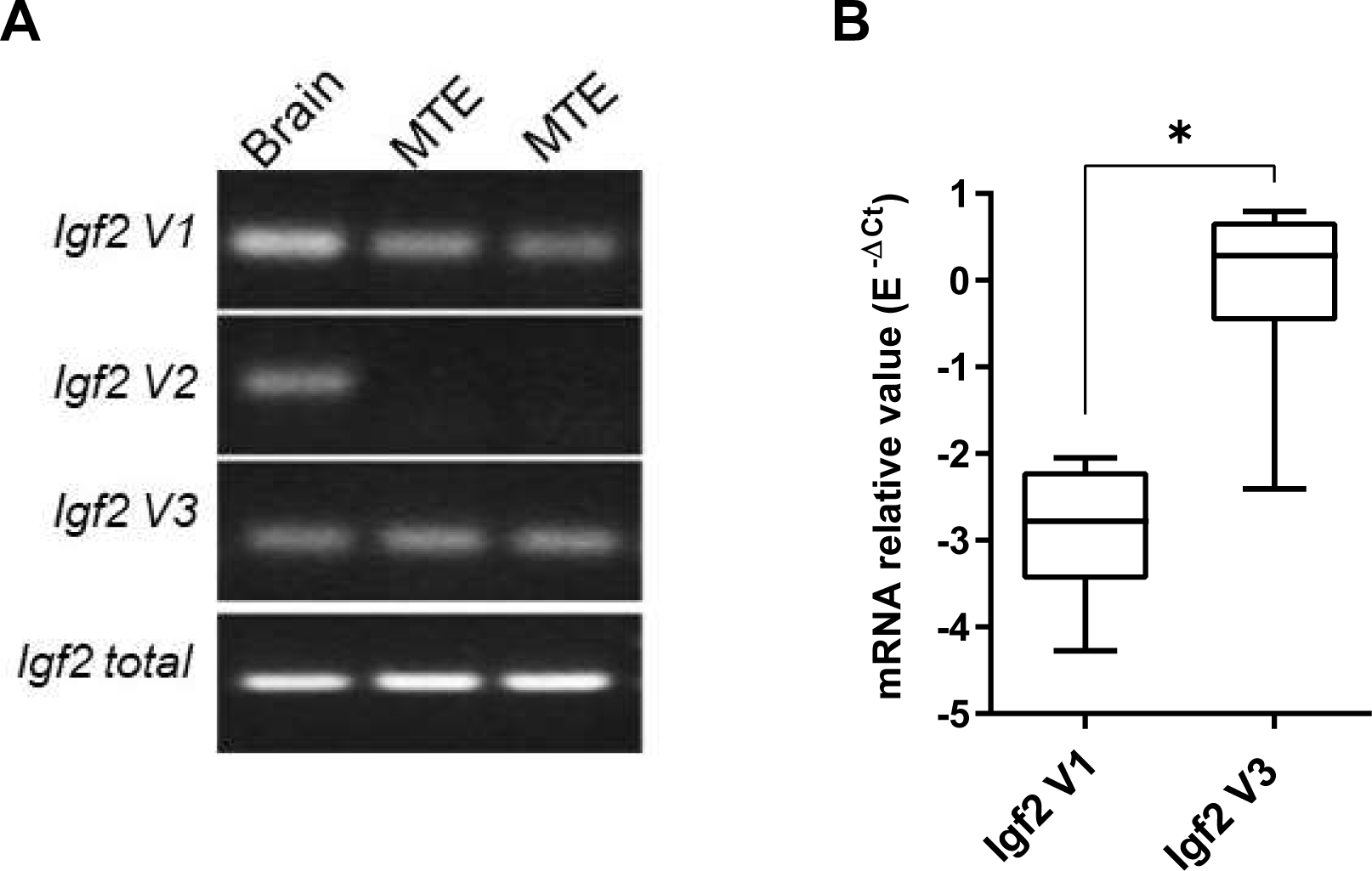
*Igf2* mRNA isoforms expression in MTE4-14 cell line. **(A)** RT-PCR of *Igf2 V1* (90 bp), *Igf2 V2* (98 bp), *Igf2 V3* (68 bp), *total Igf2* (107 bp). Mock sample from 2 independent experiments are represented, murine brain is used as positive control. **(B)** Relative mRNA expression of *Igf2 V1* and *Igf2 V3* (*n =* 6); relative expression was normalized to *Hprt* with E^−ΔCt^ formula (with E represents efficiency amplification for *Igf2 V3* and *Igf2 V1);* box-and-whisker plots extend from minimum to maximum values with lines at medians. Student *t* test, *\*p < 0.05*.

**Fig. 3.**
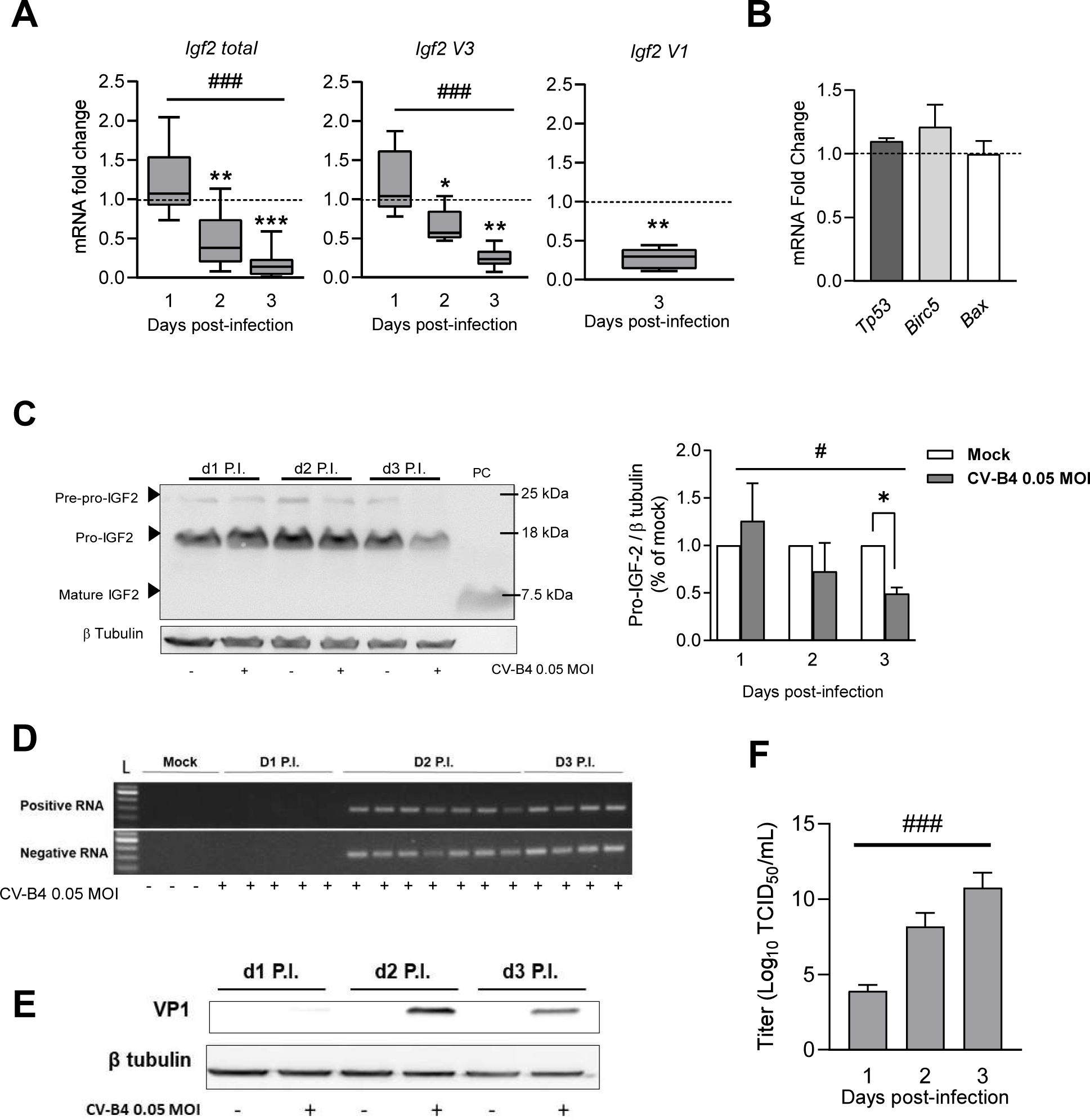
Effect of CV-B4 E2 on *Igf2* at mRNA and its precursors on MTE4-14. **(A)** Fold change of mRNA expression of *Igf2* transcripts in CV-B4 E2 _MOI = 0.05_ infected cells relative to matched mock uninfected cells (*n =* 6-12); mock samples are represented as a dashed line set at y = 1; box-and-whisker plots (CV-B4 E2 _MOI = 0.05_ infected cells) extend from minimum to maximum values, with lines at medians. **(B)** Fold change mRNA expression of *Tp53, Birc5 and Bax* in CV-B4 E2 _MOI = 0.05_ infected cells relative to matched mock uninfected cells at day 3 P.I. Mock samples are represented as a dashed line set at y = 1; data are represented as mean of fold change ± SEM; *n =* 3 **(C)** Left panel, Western blot analysis of IGF2 and its precursors in CV-B4 E2 infected cells (+) and in mock uninfected cells (−). PC, purified mature IGF2. Right panel, relative quantification of pro-IGF2 in CV-B4 E2 _MOI = 0.05_ infected cells (grey histograms) normalized to matched mock uninfected cells (white histograms) (*n =* 3); histogram represents mean of fold change ± SEM. **(D)** Representative agarose gel electrophoresis of amplicons specific to the positive and negative strands of CV-B4 E2 genome (155 bp), amplified by semi-nested strand specific RT-PCR, from CV-B4 E2 _MOI = 0.05_ infected MTE4-14 cells; mock samples served as negative control. **(E)** Representative Western blot analysis of VP1 in CV-B4 E2 _MOI = 0.05_ infected MTE4-14 cells (+) or mock uninfected cells (−). Data are representative of three independent experiments. **(F)** Viral titer of CV-B4 E2 _MOI = 0.05_ in MTE4-14 infected cells. Mean of TCID_50_/mL ± SEM (*n = 3*-5) are shown. **(A-C)** ratio paired *t* test, \*\*\**p* < 0.001 and \**p* < 0.05; **(A, C)** one-way ANOVA, #*p* < 0.05; **(F)** Kruskal-Wallis test, ###*p*<0.001.

Protein level of IGF2 and its precursors were investigated by Western Blot as in Fig. 1. As shown above *in vivo*, no mature IGF2 was detectable in mock and in CV-B4 _MOI = 0.05_ infected cells. As in Fig. 1F, we were able to detect mainly pro-IGF2 at a molecular weight comparable as in the thymus *in vivo*. During the time course of infection in MTE4-14, pro-IGF2 decreased gradually, especially at day 3 P.I. where a loss of 51% was measured in CV-B4 _MOI = 0.05_ infected cells (Fig. 3C).

Together these results show that CV-B4 is able to decrease both at mRNA and protein level IGF2 in MTE4-14 cell line. Of note as *Igf2* expression decreases, a concomitant increase of CV-B4 replication and production can be observed (Fig. 3D-F).

### CV-B4 E2 decreases *Igf2* P3 promoter activity

We hypothesized that the decrease of *Igf2 V3* mRNA (dominant *Igf2* transcript isoform in MTE4-14 cell line) in CV-B4 E2 infected cells could have a transcriptional origin. For this, we cloned *Igf2* P3 promoter (−168 to +175 relative to the TSS) upstream of Nanoluciferase coding sequence in a reporter vector (Fig. 4A). Whereas *Igf2* P3 promoter activity is unchanged at day 1 P.I., results show a significant decrease of *Igf2* P3 promoter activity at 2 days P.I. of 38% in CV-B4 _MOI = 0.05_ infected cells. Moreover, this effect increases as the CV-B4 multiplicity of infection increases (Fig. 4B). To further determine whether the *Igf2* decrease in infected cells has a post-transcriptional origin, we analyzed by RT-qPCR *Igf2 V3* mRNA expression followed by actinomycin D treatment at 2 days P.I. However, this analysis did not reveal a significant difference of *Igf2 V3* mRNA stability between mock and CV-B4 infected cells even after 10 hours of treatment; mRNA half-life was estimated at 8.6 h and 9.5 h respectively in mock and in infected cells (Fig. 4C). Together these results exclude the possibility that CV-B4 plays a significant role at the post-transcriptional level on *Igf2* expression and indicate that *Igf2* mRNA decrease is rather associated with a decrease of *Igf2* P3 promoter activity.

**Fig. 4.**
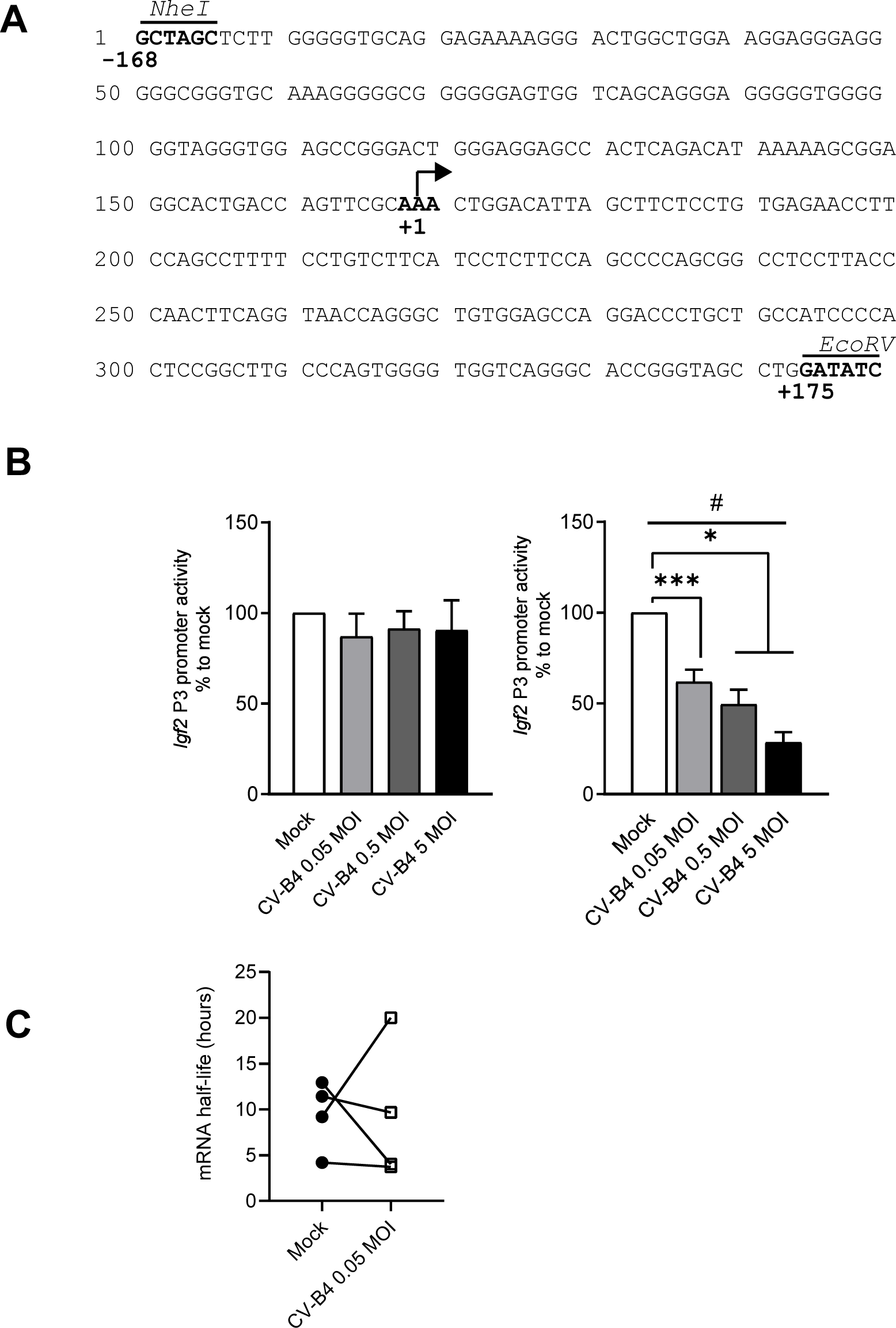
Effect of CV-B4 E2 on *Igf2* P3 promoter activity in MTE4-14 and on *Igf2* transcript stability. **(A)** Sequence of the murine *Igf2 P3* promoter sequence (−168/+175) https://epd.epfl.ch. The restriction site *NheI* and *EcoRV* are indicated above in italic. The transcription start site is represented by an arrow at +1. **(B)** Nanoluciferase relative activity of *Igf2 P3* promoter (−168/+175) after 1 (left panel) and 2 days P.I. (right panel). Analysis was realized as described in methods. Mean of relative dual-luciferase activity normalized to mock are represented ± SEM. **(C)** mRNA half-life of *Igf2 V3* transcripts in CV-B4 E2 _MOI = 0.05_ or mock uninfected cells at day 2 P.I. followed by 2-10 hours of treatment with actinomycin D (*n =* 4), vehicle control was used for data normalization for each time point. **(B-C)** ratio paired *t* test, \*\*\**p* < 0.001 and \**p* < 0.05; **(B)** one-way ANOVA, #*p* < 0.05.

Proximal promoter contains multiple binding sites specific for diverse transcription factors. To identify binding sites of *Igf2* P3 (−168 to +175 relative to the TSS) that play a role in the decrease of *Igf2* P3 promoter activity, we created a series of truncation constructs and tested them at day 2 P.I. in a transient luciferase reporter system as above (Fig. 5A). In mock and in CV-B4 _MOI = 0.05_ infected cells, the progressive deletion in *Igf2* P3 (−168 to +175) induced a gradual decrease of promoter activity, until the construct P197 (−22 to +175) where no promoter activity was detectable in all conditions (Fig. 5B). Similar results were obtained with constructs containing *Igf2* P3 promoter containing sequence downstream −22 (data not shown), identifying the region −168 to −22 as essential for *Igf2* P3 promoter minimal activity. Promoter activity of the construct P243 (−68 to +175) and P230 (−55 to +175) were significantly decreased in infected cells (Fig. 5C), while construct P291 (−116 to +175) did not reveal any significant change in infected cells indicating that the region −68 to −22 is downregulated by CV-B4 _MOI = 0.05_. To explore if the region −168 to −116 plays a role in the decrease of *Igf2* P3 promoter activity, we realized the construction P248* containing the region −168 to −116 but not the region −116 to −22. Although this region represents less than 10% of *Igf2 P3* promoter activity, we were able to detect a significant decrease of *Igf2* P3 promoter activity with the construct P248* in infected cells revealing that the region −168 to −116 is also downregulated by CV-B4 _MOI = 0.05_. We noticed that in the presence of the construct P307 (which do not contain the region −151 to −116) promoter activity in mock cells was significantly higher (*p* = 0.0242) than with the full *Igf2* P3 promoter (−168 to −116) revealing presence of potential negative regulator element in the region −151 to −116. However, results show that this element in this region does not play a role in infected cells compared to the full promoter (Fig. 5C).

**Fig. 5.**
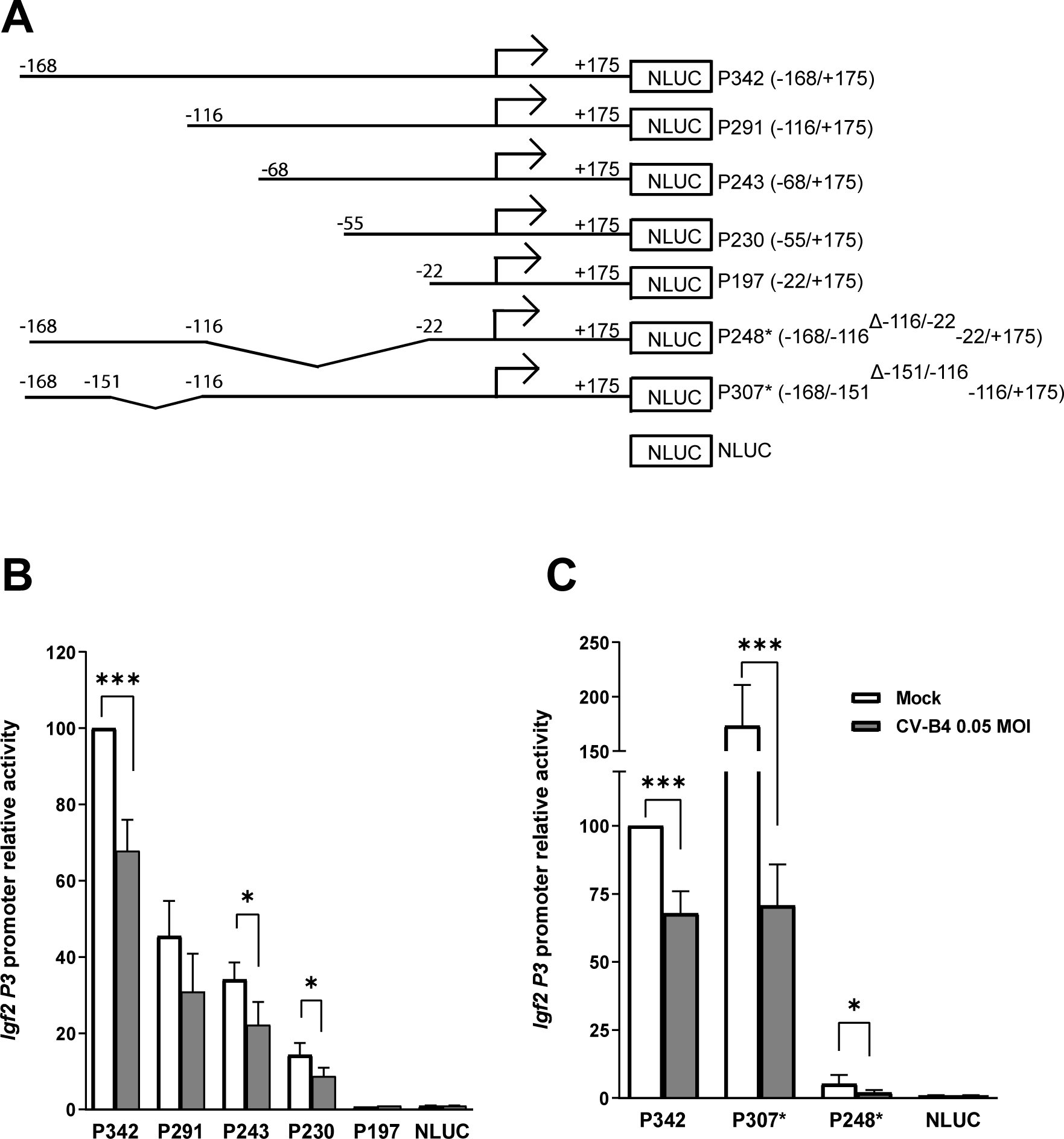
Deletion analysis of *Igf2* P3 promoter in MTE4-14 cells infected with CV-B4 E2 _MOI = 0.05._ **(A)** Schematic representation of the *Igf2* P3 promoter constructs in Nanoluciferase vector. **(B-C)** Nanoluciferase relative activity of *Igf2 P3* promoter constructs at day 2 P.I. in CV-B4 E2 _MOI = 0.05_ or in mock uninfected cells, transfected by the indicated *Igf2* P3 promoter construct. Mean of relative dual-luciferase activity normalized to mock are represented ± SEM. Mock uninfected cells transfected by the full *Igf2* P3 promoter (−168/+175) is set at 100% in each experiment (*n =* 4-16). Ratio paired *t* test, \**p* < 0.05, \*\*\**p* < 0.001.

We used then bioinformatics prediction software (21) to search for corresponding transcription factor which could bind to the region −68 to −22, representing the main region affected by CV-B4. Several putative binding sites for transcription factors were identified in this region (Table 1), as one binding site for the transcription factor Specificity Protein (SP) family, one for Krüppel-Like Factors family (KLFs), and one binding site for Zinc Finger Protein 263 (ZFP263). Binding sites for ZFP263, SP family were also identified in the region −168 to −116 (Table 1).

**Table 1.** Putative transcription factor binding site in the region −68/−22 and −168/−116.

| *Murine Igf2 P3 (-68 to -22)* |  |  |  |
| --- | --- | --- | --- |
| Name | Motif ID | GeneID | Score |
| Klf6 | M08857_2.00 | ENSMUSG00000000078 | 16.87 |
| Klf4 | M08857_2.00 | ENSMUSG00000003032 | 16.87 |
| Klf5 | M08857_2.00 | ENSMUSG00000005148 | 16.87 |
| Klf7 | M08857_2.00 | ENSMUSG00000025959 | 16.87 |
| Klf3 | M08857_2.00 | ENSMUSG00000029178 | 16.87 |
| Klf1 | M08857_2.00 | ENSMUSG00000054191 | 16.87 |
| Klf2 | M08857_2.00 | ENSMUSG00000055148 | 16.87 |
| Klf12 | M08857_2.00 | ENSMUSG00000072294 | 16.87 |
| Klf15 | M08907_2.00 | ENSMUSG00000030087 | 16.49 |
| Patz1 | M08864_2.00 | ENSMUSG00000020453 | 15.798 |
| Klf8 | M08871_2.00 | ENSMUSG00000041649 | 14.828 |
| Sp1 | M09016_2.00 | ENSMUSG00000001280 | 16.03 |
| Sp4 | M09016_2.00 | ENSMUSG00000025323 | 16.03 |
| Sp3 | M09016_2.00 | ENSMUSG00000027109 | 16.03 |
| Sp6 | M09016_2.00 | ENSMUSG00000038560 | 16.03 |
| Sp8 | M09016_2.00 | ENSMUSG00000048562 | 16.03 |
| Sp9 | M09016_2.00 | ENSMUSG00000068859 | 16.03 |
| Sp5 | M09016_2.00 | ENSMUSG00000075304 | 16.03 |
| Zfp341 | M08892_2.00 | ENSMUSG00000059842 | 15.256 |
| Zfp263 | M08272_2.00 | ENSMUSG00000022529 | 14.357 |
| Q8K439_MOUSE | M08272_2.00 | Q8K439_MOUSE | 14.357 |
| Nkx2-1 | M08136_2.00 | ENSMUSG00000001496 | 13.307 |
| Nkx2-5 | M08136_2.00 | ENSMUSG00000015579 | 13.307 |
| Nkx2-2 | M08136_2.00 | ENSMUSG00000027434 | 13.307 |
| Nkx2-6 | M08136_2.00 | ENSMUSG00000044186 | 13.307 |
| Nkx2-3 | M08136_2.00 | ENSMUSG00000044220 | 13.307 |
| Nkx2-4 | M08136_2.00 | ENSMUSG00000054160 | 13.307 |
| Nkx2-9 | M08136_2.00 | ENSMUSG00000058669 | 13.307 |

**Murine *Igf2 P3* (-168 to -116)**
| **Name** | **Motif ID** | **GeneID** | **Score** |
| --- | --- | --- | --- |
| Zfp263 | M08082_2.00 | ENSMUSG00000022529 | 14.251 |
| Q8K439_MOUSE | M08082_2.00 | Q8K439_MOUSE | 14.251 |
| Maz | M08988_2.00 | ENSMUSG00000030678 | 13.087 |
| Prdm5 | M08986_2.00 | ENSMUSG00000029913 | 13.783 |
| Zfp383 | M07689_2.00 | ENSMUSG00000099689 | 13.452 |
| Sp1 | M09016_2.00 | ENSMUSG00000001280 | 15.157 |
| Sp4 | M09016_2.00 | ENSMUSG00000025323 | 15.157 |
| Sp3 | M09016_2.00 | ENSMUSG00000027109 | 15.157 |
| Sp6 | M09016_2.00 | ENSMUSG00000038560 | 15.157 |
| Sp8 | M09016_2.00 | ENSMUSG00000048562 | 15.157 |
| Sp9 | M09016_2.00 | ENSMUSG00000068859 | 15.157 |
| Sp5 | M09016_2.00 | ENSMUSG00000075304 | 15.157 |
| Etv3 | M09055_2.00 | ENSMUSG00000003382 | 13.617 |
| Etv1 | M09055_2.00 | ENSMUSG00000004151 | 13.617 |
| Etv2 | M09055_2.00 | ENSMUSG00000006311 | 13.617 |
| Elk3 | M09055_2.00 | ENSMUSG00000008398 | 13.617 |
| Gabpa | M09055_2.00 | ENSMUSG00000008976 | 13.617 |
| Elk1 | M09055_2.00 | ENSMUSG00000009406 | 13.617 |
| Etv5 | M09055_2.00 | ENSMUSG00000013089 | 13.617 |
| Fli1 | M09055_2.00 | ENSMUSG00000016087 | 13.617 |
| Etv4 | M09055_2.00 | ENSMUSG00000017724 | 13.617 |
| Ets2 | M09055_2.00 | ENSMUSG00000022895 | 13.617 |
| Elk4 | M09055_2.00 | ENSMUSG00000026436 | 13.617 |
| Elf4 | M09055_2.00 | ENSMUSG00000031103 | 13.617 |
| Ets1 | M09055_2.00 | ENSMUSG00000032035 | 13.617 |
| Elf1 | M09055_2.00 | ENSMUSG00000036461 | 13.617 |
| Elf2 | M09055_2.00 | ENSMUSG00000037174 | 13.617 |
| Erg | M09055_2.00 | ENSMUSG00000040732 | 13.617 |
| Erf | M09055_2.00 | ENSMUSG00000040857 | 13.617 |
| Fev | M09055_2.00 | ENSMUSG00000055197 | 13.617 |
| XP_911724.4 | M09055_2.00 | XP_911724.4 | 13.617 |
| Zfp263 | M07844_2.00 | ENSMUSG00000022529 | 16.224 |
| Q8K439_MOUSE | M07844_2.00 | Q8K439_MOUSE | 16.224 |
| Zfp281 | M08999_2.00 | ENSMUSG00000041483 | 13.023 |
| E2f4 | M09034_2.00 | ENSMUSG00000014859 | 14.084 |
| E2f3 | M09034_2.00 | ENSMUSG00000016477 | 14.084 |
| E2f2 | M09034_2.00 | ENSMUSG00000018983 | 14.084 |
| E2f1 | M09034_2.00 | ENSMUSG00000027490 | 14.084 |
| E2f5 | M09034_2.00 | ENSMUSG00000027552 | 14.084 |
| E2f6 | M09034_2.00 | ENSMUSG00000057469 | 14.084 |
| Sall4 | M08866_2.00 | ENSMUSG00000027547 | 13.786 |
| Patz1 | M08287_2.00 | ENSMUSG00000020453 | 14.129 |

### CV-B4 E2 alters IL-6/STAT3 signaling and decreases STAT3 phosphorylation

Given that Signal transducer and activator of transcription 3 (STAT3) acts as positive regulators of *Igf2* transcripts in human and in mice (22, 23), STAT3 protein expression and phosphorylation (STAT3 ^pY705^) level were analyzed by Western Blot. Results show that CV-B4 _MOI = 0.05_ decreases gradually STAT3^pY705^ during the course of the infection, whereas STAT3^total^ was not significantly affected. STAT3^pY705^ was decreased of 51% and 65% respectively at 2 and 3 days P.I. (Fig. 6A). In line with this result, *Bcl2,* a STAT3 response gene (24), was similarly decreased during the time course of CV-B4 infection in MTE4-14 (Fig. 6B).

**Fig. 6.**
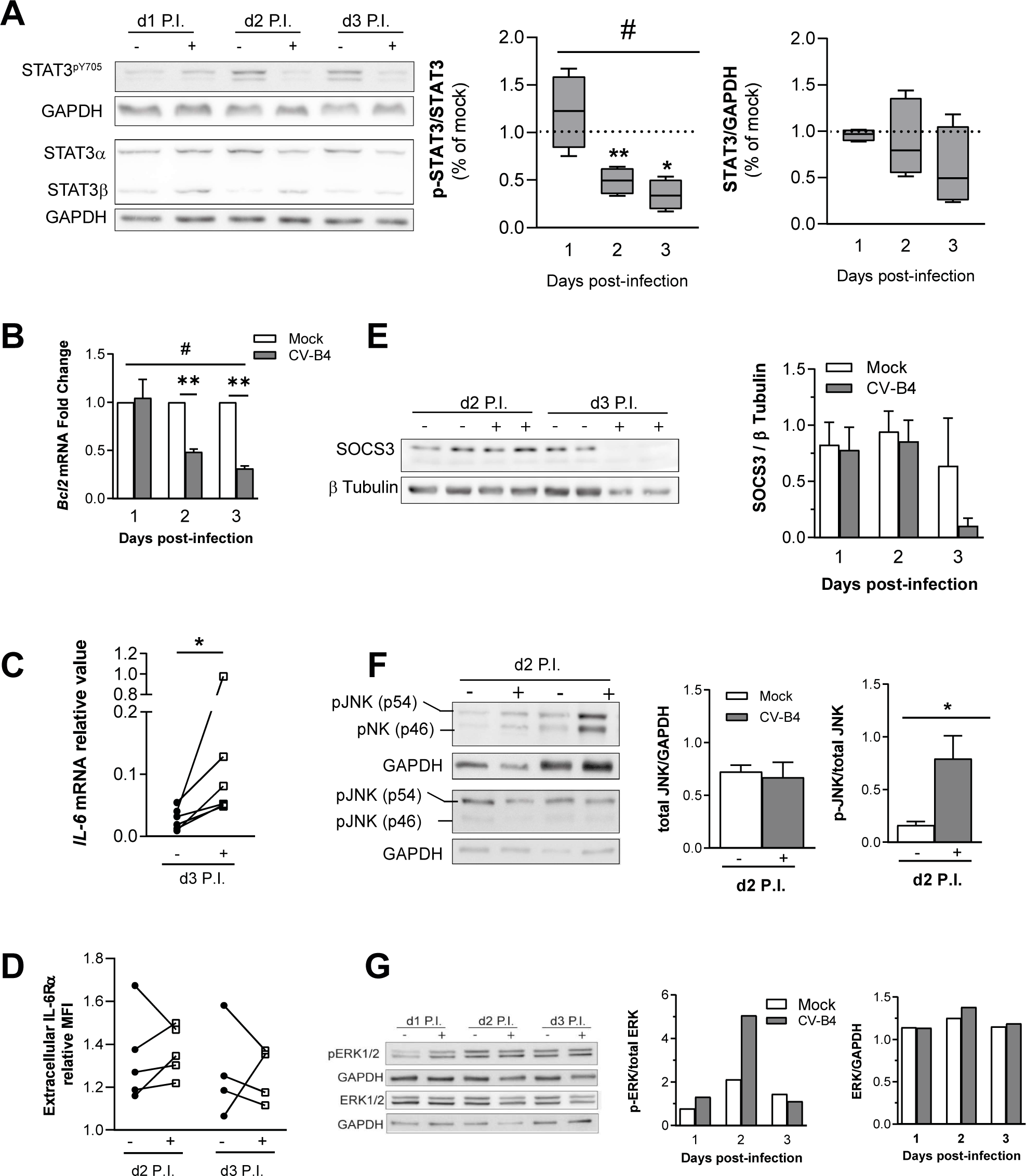
Effect of CV-B4 E2 on STAT3 phosphorylation and on STAT3 signaling pathway. **(A)** Left panel, Western blot analysis of STAT3^total^ and STAT3^pY705^ in CV-B4 E2 infected cells (+) and in matched mock uninfected cells (−). Right panel, relative quantification of STAT3^total^ and STAT3^pY705^ in CV-B4 E2 _MOI = 0.05_ infected cells (+) and in day matched mock uninfected cells (−) (*n =* 4-5); box-and-whisker plots representing CV-B4 E2 _MOI = 0.05_ infected cells, extend from minimum to maximum values, with lines at medians; mock samples are represented as a dashed line set at y = 1. **(B)** Fold change of *Bcl2* mRNA in CV-B4 E2 _MOI = 0.05_ infected cells (grey histograms) relative to matched mock uninfected cells (white histograms); n = 3-6. **(C)** *Il6* mRNA relative expression in CV-B4 E2 _MOI = 0.05_ infected cells (+) and in matched mock uninfected cells (−); *n =* 6. **(D)** Quantification of extracellular IL-6Rα by flow cytometry in CV-B4 E2 _MOI = 0.05_ infected cells (+) and in matched mock uninfected cells (−); data are represented as MFI value relative to isotype control; *n = 4*-5. **(E)** Left panel, Western blot analysis of SOCS3 in CV-B4 E2 _MOI = 0.05_ infected cells (+) in day matched mock uninfected cells (−). Right panel, relative quantification of SOCS3 in CV-B4 E2 _MOI = 0.05_ infected cells (+) and in day matched mock uninfected cells (−); *n = 2*-3. **(F)** Left panel, Western blot analysis of JNK ^total^ and phosphorylated JNK in CV-B4 E2 _MOI = 0.05_ infected cells (+) and in matched mock uninfected cells (−); two independent experiments are represented. Right panel, relative quantification of JNK ^total^ and phosphorylated JNK in CV-B4 E2 _MOI = 0.05_ infected cells (+) and in matched mock uninfected cells (−); *n =* 3. **(G)** Left panel, Western blot analysis of total ERK and p-ERK in CV-B4 E2 _MOI = 0.05_ infected cells (+) and in matched mock uninfected cells (−). Right panel, relative quantification of ERK^total^ and phosphorylated ERK in CV-B4 E2 _MOI = 0.05_ infected cells (+) and in matched mock uninfected cells (−); *n =* 1. **(B, E, F-G)** Histograms represents average ± SEM. **(A-F)**, Ratio paired *t* test, \*\**p* < 0.01 and \**p* < 0.05; **(A-B)** one-way ANOVA, #*p* < 0.05.

However, *Il6*, activator of *STAT3* ^pY705^, was upregulated in infected cells (Fig. 6C) which is in discordance with the observed decrease of STAT3^pY705^. We then investigated expression of the receptor IL6Rα (CD126) by flow cytometry but we were not able to detect a significant difference between mock and infected cells (Fig. 6D).

Previous reports indicate that suppressor of cytokine signaling 3 (SOCS3), a negative regulator of STAT3 activation, is induced by CV-B3 (25, 26). SOCS3 protein expression was analyzed by Western Blot and was not upregulated during the infection. On the opposite, SOCS3 protein level tends to decrease during the infection (Fig. 6E). More recently shown to play a negative role in STAT3^pY705^ activation (27–30), expression of the c-Jun N-terminal Kinase (JNK) and the Extracellular signal-Regulated protein Kinase (ERK) were analyzed in MTE4-14 infected cells. At 2 days P.I., we observed that phosphorylated JNK and phosphorylated ERK were both increased whereas level of JNK^total^ or ERK^total^ remained unchanged (Fig. 6F-G).

## DISCUSSION

In this research, we firstly investigate the effect of CV-B4 E2 on *Igf2* expression in murine TECs *in vivo*. The analysis of *Igf2* transcripts isoforms in mock inoculated mice revealed that *Igf2 V3* and *Igf2 V1* are the isoforms which can be mainly detected in CD45^−^ enriched TECs. These results corroborate with previous reports on human TECs showing that *IGF2* P3 and *IGF*2 P4, homologous promoter of *Igf2 V1* and *Igf2 V3* respectively are active in these cells (31) and accordingly, much less *Igf2 V2* in CD45^−^ enriched TECs was detected. In CV-B4 inoculated mice, both *Igf2 V3* and *Igf2 V2* were decreased followed by an upregulation of *Igf2 V1*. These results explained why *total Igf2* (representing quantitatively sum of all *Igf2* isoforms) was not decreased in infected mice and indicate a differential regulation of *Igf2* mRNA transcripts in the thymus by CV-B4 E2. Thus, distinct pathways may regulate *Igf2 V3* and *V1* mRNA isoforms during CV-B4 infection. This temporal decrease of *Igf2 V3* and *Igf2 V2* was nevertheless sufficient to achieve a later significant decrease of Pro-IGF2. Consequently a role of CV-B4 on IGF2 presentation in the thymus cannot be excluded, indeed it was shown that CV-B3 is able to alter class I antigen presentation (32). Nevertheless, we were not able to detect any mature IGF2 neither in the thymus nor in the TEC cell line MTE4-14. A differential processing of IGF2 may occur in the thymus. It was reported that in rat, mature forms of IGF2 are also not found in the brain (33). Moreover, sequencing data of TECs indicates low mRNA expression level of protein convertase PC4 (*Pcsk4*), enzyme cleaving pro-IGF2 in mature IGF2. A low thymic expression of *Pcsk4* would affect IGF2 processing and would allow only the detection of IGF2 immature precursors (34, 35). In mice inoculated with CV-B4, we were able to detect CV-B4 RNA in the pancreas. However, CV-B4 RNA was not detected in the thymus which is in discordance with previous reports reporting the presence of CV-B4 RNA in the whole thymus (9, 36). It is likely that the method we employed here for TECs enrichment has interfered negatively with the detection of CV-B4 as it only allows to detect CV-B4 RNA within the cells (Table 2). Nonetheless, this result indicates that CV-B4 could not replicate as much as expected within murine thymic cells. Of note, attempts to detect CV-B3 replication within cells of murine thymus have similarly failed (37, 38).

**Table 2.** CV-B4 E2 mRNA detection by RT-PCR performed in other studies.

| Tissue | Days P.I. |  |  |  |  | Study | Processing | Viral dose<br>(TCID<sub>50</sub>) | Inoculation<br>method |
| --- | --- | --- | --- | --- | --- | --- | --- | --- | --- |
|  | 2 | 3 | 7 | 8 | 10 |  |  |  |  |
| Pancreas | 2/2 | 2/2 | 2/2 | n/a | 2/2 | *Jaïdane et. al*, [2006] | WT | 10<sup>4.74</sup> | Oral |
|  | n/a | n/a | n/a | 2/2 | n/a | *Aguech-Oueslati et. al*,<br>[2017] | WT | 10<sup>5</sup> | Oral<br>I.P. |
|  | 6/6 | 6/6 | 2/6 | n/a | n/a | Actual study | WT | 10<sup>6</sup> | I.P. |
| Thymus | 2/2 | 2/2 | 2/2 | n/a | 1/2 | *Jaïdane et. al*, [2006] | WT | 10<sup>4.74</sup> | Oral |
|  | n/a | n/a | n/a | 2/2 | n/a | *Aguech-Oueslati et. al*,<br>[2017] | WT | 10<sup>5</sup> | Oral + I.P. |
|  | 0/6 | 0/6 | 0/6 | n/a | n/a | Actual study | EP | 10<sup>6</sup> | I.P. |

WT, whole tissue; EP, enzymatic processing; I.P., intra-peritoneal; n/a, no data; 2/2 represents two positive samples for two independent biological samples tested

Interestingly, several reports show that interferon (IFN)-α or IFN-β are able to decrease *Igf2* expression (39–41) and that inactivated CV-Bs virions, by interacting with extracellular Toll-like receptors (TLRs), can still upregulate these interferon suggesting that no replication within cells is actually required to induce IFN expression (42). Thus, CV-B4 draining not replicative virions, within the thymus could induce via TLRs IFN-α or IFN-β which may then alter *Igf2* transcripts.

The second objective of this study was to explore mechanisms of *Igf2* decrease in MTE4-14 cells infected with CV-B4 E2. Similar to the results obtained *in vivo, Igf2 V3* and *Igf2 V1* are both detected in the cell line with a dominance of *Igf2 V3* reflecting a dominant activity of *Igf2 P3* and a lower activity of *Igf2 P2.* Both *Igf2 V3* and *Igf2 V1* were downregulated concomitantly to the increase of CV-B4 replication and production. Given that *Igf2 V1* was on the opposite increased *in vivo* and CV-B4 E2 viral RNA not replicative (or under the limit of detection) within murine thymic cells, a differential regulation scenario of *Igf2 V1* isoforms, possibly related to CV-B4 replication, might takes place in the MTE4-14 cell line, where a higher rate of CV-B4 replication occurred.

In this study, we demonstrate that the decrease of *Igf2 V3* major isoforms is due to a decrease of *Igf2 P3* promoter activity (−168 to +175 relative to the TSS) revealing a downregulation of *Igf2 V3* at the transcriptional level. *Igf2 V3* mRNA stability was not affected in infected cells which confirm the regulation of *Igf2 V3* mRNA at the transcriptional level. Previous studies on *total Igf2* mRNA stability indicate that *total Igf2* is a highly stable transcript (43, 44). Our results demonstrate also the promoter region of *Igf2 P3* −68 to −22 (relative to the TSS) and secondary the region *Igf2 P3* −168 to −116 (relative to the TSS) as regulated negatively by CV-B4. Nonetheless, as we used only CV-B4 _MOI = 0.05_ for this analysis, we can expect that with higher MOI, others *Igf2 P3* promoter regions might be affected. Predictive analysis of *Igf2 P3* −68 to −22 (relative to the TSS) shows binding sites corresponding to transcription factor from the SP/KLF family. Among them, SP1 was accordingly identified as positive activator of human *Igf2 P4* (homologous to the murine *Igf2 P3* promoter) (45). Besides this, the SP/KLFs transcription factor families were shown to play essential roles in differentiation, development, proliferation or cell death regulation. Interestingly, KLFs transcription factors play a role for thymocytes development (46–48). Further studies would be required to explore *Igf2* regulation by KLFs during CV-B4 infection which could link the observed alteration of thymocytes differentiation (10, 11) and the decrease of *Igf2 V3*.

STAT3, playing a role in *Igf2* expression, has been shown recently to be indispensable for TECs development and survivals (49–51). Here in this study, we identified also that CV-B4 decreases STAT3^pY705^ in a thymic epithelial cell line. Moreover, the analysis of the *Igf2 P3* region −168 to −116 (related to the TSS) revealed an E2F binding site which may be recognized by STAT3 (52). It is then possible that the decrease of *Igf2 P3* promoter activity would be linked, at least, in part to a decrease of STAT3^pY705^. Additionally, as *Il6* was upregulated in infected cells, this work revealed an inhibition of IL-6/STAT3 signaling. This upregulation has been previously reported in TECs and in MTE4-14 as well (12, 14). STAT3^pY705^ inhibition was neither not related to an upregulation of SOCS3, nor by an inhibition of IL-6Rα (CD126). On the contrary, we observed a decrease of SOCS3 which can be attributed to the decrease of STAT3 ^pY705^ signaling itself. Our results indicate an upregulation of JNK and ERK phosphorylation. These kinases, important for enterovirus viral progeny release (53–55) can also indirectly contribute to STAT3^pY705^ downregulation (27–30), which suggest that induction of phosphorylated kinases JNK and ERK could contribute to STAT3^pY705^ decrease and potentially plays also a role in the decrease of *Igf2* in MTE4-14 cells infected with CV-B4.

Together, these findings bring new knowledge of *Igf2* regulation by CV-B4 in the context of a thymic infection, and further support the idea that CV-B4 may disturb *Igf2* thymic presentation and decrease central tolerance to insulin.

## METHODS

### Cells and virus

The murine thymic epithelial cell line MTE4-14, of medullar origin, derived from C3H/J (*H-2^k^*) thymic neonatal lobes (13), was grown in complete high glucose Dulbecco’s modified Eagle medium (DMEM; Gibco) supplemented with 10% heat-inactivated fetal calf serum (FCS; Gibco), 2 mM L-glutamine (Gibco), 0.1 μg/mL epidermal growth factor (EGF; Sigma-Aldrich), 100 U/mL penicillin and 100 μg/mL streptomycin (Gibco) (13, 14). VERO cells (kindly provided by the Laboratory of Virology and Immunology, Giga, university of Liège) were grown in DMEM high glucose supplemented with 10% FCS, 100 U/mL penicillin and 100 μg/mL streptomycin. CV-B4 E2, the diabetogenic strain of coxsackievirus B4, (kindly provided by Ji-Won Yoon, Julia McFarlane Diabetes Research Center, Calgary, Alberta, Canada) was propagated in Hela cell line in DMEM high glucose supplemented with 10% FCS. Supernatants were collected 4 days after inoculation, after three freeze/thaw cycles then clarified by centrifugation at 2,500 × g for 10 min, filtered and stored at 80 °C.

### Mice inoculation and infection by CV-B4 E2

For *in vivo* experiment, 4–6 weeks old Swiss Albino female mice were used (Janvier Laboratories). Mice were inoculated by intraperitoneal route with either 100 μL of sterile DPBS (DPBS, Dulbecco’s Phosphate-Buffered Saline, Thermo Fisher Scientific) for mock-infected mice or with 100 μL of CV-B4 E2 diluted in sterile DPBS at 1.10^6^ TCID_50_/mL. Mice were grouped with a maximum of six per cage, checked and weighted daily. Mice were treated according to general ethic rules with unlimited access to food and water. Six infected and mock-infected animals were sacrificed 2, 3 and 7 days P.I. From each animal, thymus and pancreas were collected. All procedures were approved by the university hospital of Liège ethics committee (Protocol n°13-1611).

### TEC isolation, enrichment and immunostaining

Protocol was carried out as previously described (16). Briefly, thymus lobes were cut in small pieces and cleaned for blood and connective tissue in RPMI (Lonza) supplemented with 10% FCS, 2 mM L-glutamine and 100 U/mL penicillin and 100 μg/mL streptomycin (Gibco). Thymic fragments were washed in RPMI 2% FCS and digested 15 min at 37 °C in 500 μg/mL Liberase TL (Sigma-Aldrich) and 111 μg/mL DNase I from bovine pancreas (Sigma-Aldrich). Thymic fragments were mixed in the beginning and in the end of enzymatic digestion. Resulting supernatant was incubated 5 min on ice in DPBS supplemented with 1% FCS and 5 mM EDTA pH7.3. Complete RPMI was added on supernatant and the thymic cell suspension was centrifuged and filtered. Fifty million cells were used for TECs enrichment with mouse CD45 microbeads (Miltenyi Biotec) according to the manufacturer’s protocol. For immunostaining, CD45^−^ enriched cells (TECs) and total thymic suspension was incubated with anti-CD16/CD32 Fc block 1 50 (clone 93, eBioscience) during 15 min at 4 °C in MACS buffer followed by incubation with anti-CD45 1:100 (clone 30F-11, BD biosciences), anti-EpCAM 1:100 (clone G8.8, eBioscience) during 30 min. Samples were analyzed on a FACSCanto flow cytometer (BD biosciences) and the raw data analyzed with FlowJo software (Treestar). Purity of CD45 negative sorted cells was in average 80% (data not shown). Aliquots of total thymic cells (digested unsorted cells) were stored for protein and RNA extraction, aliquots of CD45^−^ isolated cells were stored for RNA extraction.

### Infection of MTE4-14 by CV-B4 E2

MTE4-14 cells were seeded at 150,000 cells per well on 6-well culture plates in DMEM high glucose supplemented with 10% heat-inactivated FCS, 2 mM L-glutamine, 0.1 μg/mL EGF and incubated overnight at 37 °C. The culture medium was then removed and cells were inoculated with 500 μL per well of CV-B4 E2 in DMEM only with a multiplicity of infection (MOI) of 0.05. The MOI is defined by the formula:

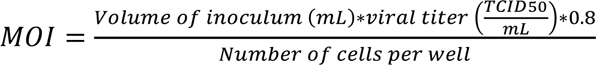

Alternatively, 2.10^4^ MTE4-14 cells were seeded on 96-well culture plates flat bottom and were inoculated with 100 μL per well with various MOI ranging from 0.05 to 5. One well served as control for cell count before infection, allowing similar MOI between experiments. Mock-infected cells served as negative control and were treated under same conditions for all experiments, except inoculation with DMEM alone. After 90 min of incubation cells were washed with DMEM alone, and incubated with complete DMEM without antibiotics for 3 days. At 1 to 3 days P.I., one plate was stopped and processed as follow: culture supernatants were removed and stored at −20 °C for CV-B4 E2 titration, cells were washed, scrapped in PBS then were processed for RNA or protein extraction.

### Modified TCID_50_ titration assay

CV-B4 E2 virus titer was determined on VERO cells by a modified Reed and Muench limiting dilution assay as previously described (56, 57). Briefly, after an incubation of 7 days with CV-B4 E2 dilutions, cells were incubated during 3 hrs with MTT reagent (Sigma-Aldrich), then formazan was dissolved in DMSO and absorbance was measured at 550 nm. TCID_50_ values (limiting dilution corresponding to 50% of viability) were obtained by the use of the V_50_ parameter of Boltzmann sigmoid function.

### Flow cytometry

For MTE4-14 flow cytometry analysis, cells were harvested with EDTA 5 mM in DPBS during 15 min at 37 °C, washed with 10% FBS before addition of Fc block as described above. MTE4-14 cells were then stained with anti-mouse CD126 APC 1:40 (D7715A7, Biolegend) or with APC rat IgG2b, κ isotype control (Biolegend).

### Reverse transcription, end point PCR and real-time quantitative PCR

Total RNA was extracted with Nucleospin RNA kit (Machery Nagel) according to manufacturer’s instructions. Alternatively, RNA was extracted with TRIzol Reagent (Ambion) following manufacturer’s instructions. If TRIzol was used, contaminating DNA was eliminated with 2 U of DNase Turbo (Ambion) during 30 min at 37 °C, followed by RNA re-extraction with phenol chloroform. RNA concentration was measured on Nanodrop ND-1000 (Thermo Fisher Scientific). A_260/280_ ratio upper than 1.8 were considered as acceptable. Reverse transcription was performed using 200-500 ng of total RNA in the Transcriptor First Strand cDNA synthesis kit (Roche) with 60 μM random hexamer and 6.25 μM oligo(dT)_18_ primer. Reverse transcriptase minus control was used as control for DNA contamination. End Point PCR were realized on an iCycler Thermal Cycler (Bio-Rad) in the presence of GoTaq G2 polymerase (Promega) in 25 μL containing 10-25 ng cDNA, 200 μM dNTPs, 200 nM of each forward and reverse primer, 25 mM MgCl_2_, 1X Green Buffer and 0.625 U of GoTaq G2 polymerase. The PCR parameters were: one cycle at 95 °C for 2 min; 35 cycles with denaturation at 95 °C for 30 s, annealing at 60 °C for 30 s and elongation at 72 °C for 30 s; one cycle at 72 °C for 5 min. PCR products were visualized on agarose gel. Real time PCR was performed using Takyon No Rox Sybr 2X masterMix blue dTTp (Eurogentec) with 200 nM of each primer on an iCycler iQ(Bio-Rad). qPCR parameters were: initial denaturation at 95 °C for 10 min; 40 cycles with denaturation at 95 °C for 10 s, annealing at 60 °C for 30 s and amplification at 72 °C for 25 s. Each qPCR reaction was ended by a melting curve with a ramp of 0.5 °C from 55 to 95 °C to control single PCR product amplification. A negative template control for each gene was included in the plate for each mix. Gene expression values were calculated based on the comparative Ct normalized to *Hprt* and displayed as fold change to mock (2^−ΔΔCt^) or in relative value (2^−ΔCt^). Primers (Table 3-1) were double checked with PRIMER-BLAST.

**Table 3-1.**
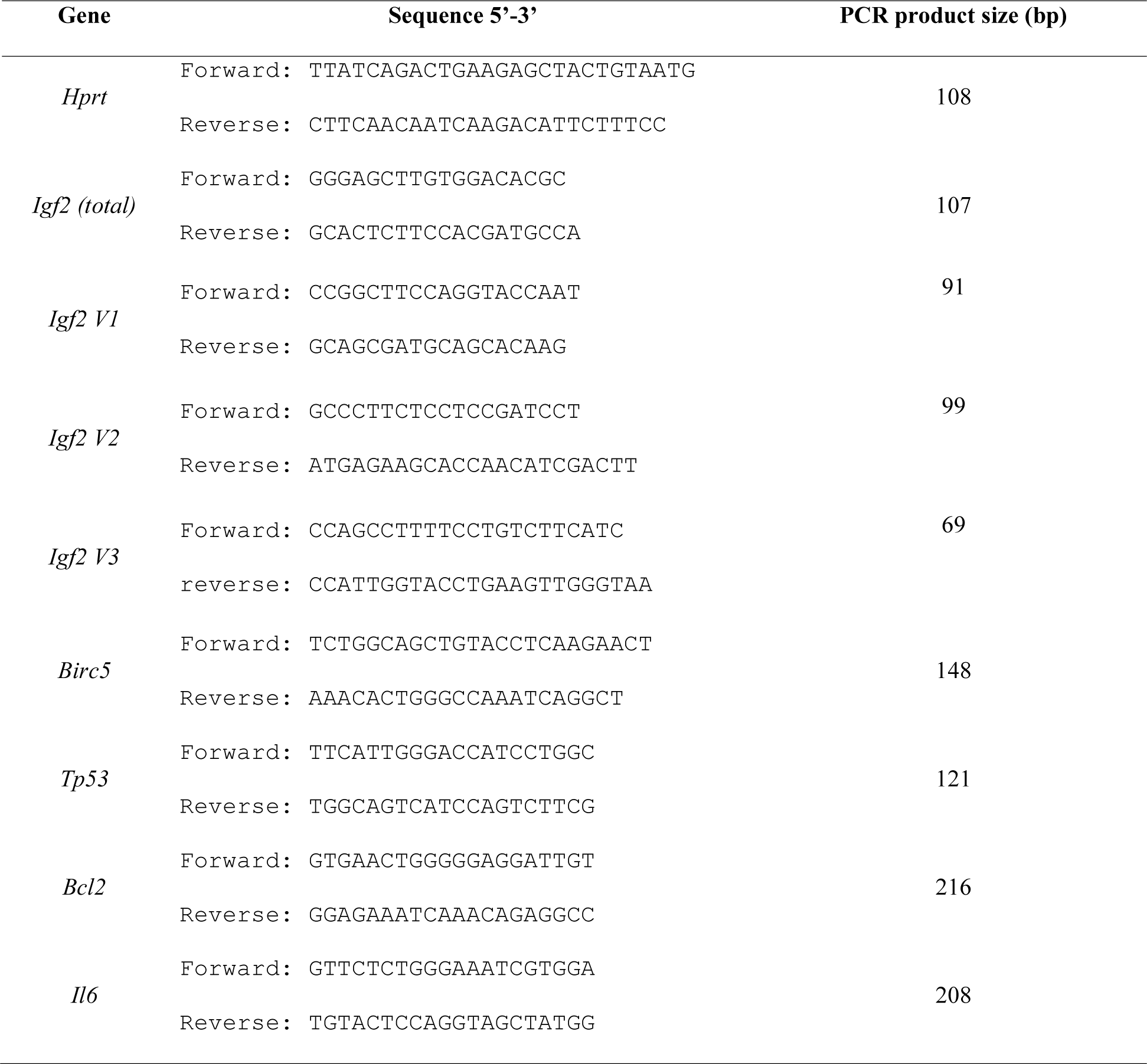
qPCR primers sequence.

### One-step CV-B4 E2 RNA detection and two-step CV-B4 E2 RNA detection by PCR

One-step RT-PCR was realized with the SuperScript III One-Step RT-PCR System with Platinum Taq DNA Polymerase (Thermo Fisher Scientific) on an iCycler Thermal Cycler (Bio-Rad). Reactions were performed following manufacturer’s instruction containing 500 ng of RNA, 100 nM of 007 and 008 primers. RT-PCR parameters were: one step 50 °C 30 min for cDNA synthesis, followed by one cycle at 94 °C 2 min for initial denaturation; 38 cycles with one cycle 94 °C for 30 s, annealing at 60 °C and extension at 68 °C for 30 s and a final extension at 72 °C for 10 min. PCR products were visualized on agarose gel (1.5%). In case of negative amplification, PCR product was run for a semi-nested PCR for 35 cycles in the presence of Gotaq G2 DNA polymerase as described above. For two-step CV-B4 E2 RNA detection, reverse transcription was performed as described above in 10 μL with 1 μM reverse primer 007 (5’-ATTGTCACCATAAGCAGCCA-3’) for positive strand of CV-B4 E2 or 1 μM forward primer 008 (5’-GAGTATCAATAAGCTGCTTG-3’) for negative strand detection of CV-B4 E2. PCR amplification was performed then with 100 nM of 007 and 008 primers. PCR parameters were: one cycle at 95 °C for 2 min; 25 cycles with denaturation at 95 °C for 30 s, annealing at 60 °C and extension at 72 °C for 30 s. PCR was ended 5 min at 72 °C. PCR products were analyzed on agarose gel and give a PCR product of 412 bp. Negative PCR was run for semi-nested PCR with internal primer 006 (5’-TCCTCCGGCCCCTGAATGCG-3’) and 007. PCR parameters were: one cycle at 95 °C for 2 min; 35 cycles with denaturation at 95 °C for 30 s, annealing at 60 °C and extension at 72 °C for 30 s. PCR was ended 5 min at 72 °C. Semi-nested PCR product size is 155 bp.

### *Igf2* P3 Nluc plasmid construction and site-directed mutagenesis

The 342 bp *Igf2* P3 promoter sequence (Fig. 4A), containing the proximal promoter of the murine *Igf2* P3 (58, 59) was synthetized by the GeneArt Gene Synthesis service (Thermo Fisher Scientific). TSS sequence was localized with EPD database (60), *Igf2 P3* promoter sequence was then cloned in the Nanoluciferase expressing plasmid pNL1.2 NlucP plasmid (Promega) by the use of restriction site *NheI* (5’) and *EcoRV* (3’) in *Igf2* P3 promoter by the GeneArt Gene Synthesis service. The resulting plasmid was called *Igf2 P3* Nluc. Site-directed mutagenesis or promoter deletions were performed with the Q5 Site-Directed Mutagenesis Kit (New England Biolabs). PCR reactions were done in 50 μL containing 25 μL of Q5 Hot Start High-Fidelity 2X Master Mix, 500 nM of each forward and reverse primer (Table 3-2) and 50 ng of plasmid. The PCR parameters were: one cycle at 98 °C for 30 s; 25 cycles with denaturation step 98 °C for 10 s, annealing (Table 3-2) for 30 s and extension step at 72 °C for 150 s and ended at 72 °C for 2 min. Three annealing temperature were performed in parallel for each primer set. PCR products were loaded on 1% agarose gel with Orange DNA Loading Dye 6X (Thermo Fisher Scientific). SYBR safe (Thermo Fisher Scientific) was added on agarose gel at 1:10,000 for DNA visualization. When single PCR product was detected, KLD mix was added on PCR reaction. Briefly, the KLD reaction contains 2.5 μL 2X KLD buffer, 1 μL of Enzyme Mix, 0.5 μL of PCR product and 0.5 μL of nuclease-free water. Five μL of KLD reaction were mixed with 25 μL of *Escherichia coli* DH5α chemocompetent strain and transformed 45 s at 42 °C. Transformed bacteria were selected on agar plate containing ampicillin at 100 μg/ml. Plasmids were extracted with NucleoSpin Plasmid Transfection-grade (Machery-Nagel) and concentration were measured on Nanodrop ND-1000 (Thermo Fisher Scientific). All plasmids were sequenced by Sanger method (Giga-Genomics).

**Table 3-2.**
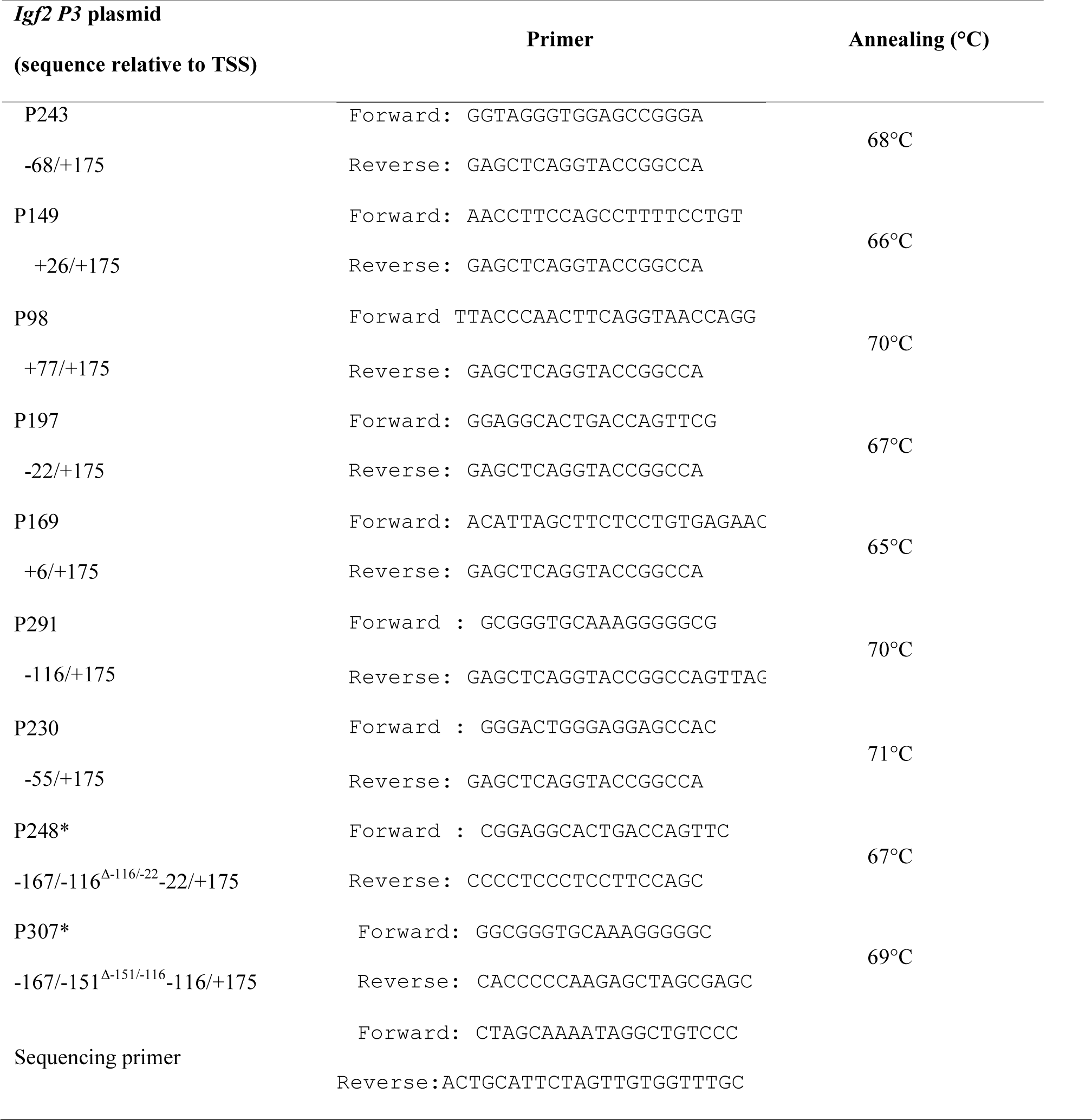
Site-directed mutagenesis primers sequence.

### Transfection and dual luciferase reporter assay on MTE4-14 cell line

Plasmid transfection was performed in 96-well plate. Briefly, each reaction contained 20 μL of 100 ng of pGL3 plasmid (Promega), mixed with 100 ng *Igf2* P3 Nluc (or empty vector or constructs obtained from *Igf2* P3 Nluc), 0.5 μL of Genius DNA transfection Reagent (Westburg) diluted in DMEM. Mix were added directly on 2.10^4^ cells/well in a 96-well plate during the seeding. After an overnight incubation at 37 °C, cells were then infected by CV-B4 E2 or DMEM only as described above. Cells were infected during a maximum of two days. Nanoluciferase activity and firefly luciferase activity, were analyzed by the Nano-Glo Dual-Luciferase Reporter Assay System (Promega) following manufacturer’s instructions. Bioluminescence was analyzed with a FilterMax F5 (Molecular Devices). Normalized luciferase activity was calculated as the ratio of the Nanoluciferase activity to the firefly luciferase for each sample, then normalized to empty vector for mock uninfected cells and for CV-B4 E2 infected cells. In each experiment, relative mock value was then set up to 100%.

### mRNA stability assay

Actinomycin D (Sigma-Aldrich) was prepared at 1 mg/mL in ethanol and used at 5 μg/mL. mRNA half-life of *Igf2 V3* was estimated by linear regression in each experiment from *Igf2 V3* relative value (2^−ΔCt^) (normalized to value of vehicle control for each time point) with the formula 2^−ΔCt(*Igf2V3-Hpr*)^ = f(time of actinomycin D treatment) for mock or CV-B4 E2 infected cells.

### SDS-PAGE and Western Blot

Proteins were extracted in RIPA buffer supplemented with protease and phosphatase inhibitor mini tablets cocktail (Pierce) and stored at −20 °C for further use. Protein extracts were quantified with BCA protein assay kit (Pierce) and denatured at 95 °C during 5 min in Laemmli buffer supplemented with β-mercaptoethanol. Ten μg (or fifty μg for IGF2 detection) of protein lysates were separated on 12% SDS-PAGE gel then transferred to PVDF membrane (Amersham). Transfer was performed at 4 °C during 90 min at 80 V in transfer buffer containing 25 mM Tris, 192 mM glycin and 20% (v/v) methanol. Prior to immunoblotting, membranes were blocked in blocking buffer (5% w/v BSA in TBS-T) during 1 hr at room temperature and primary antibodies (Table 4) diluted in blocking buffer were added for overnight at 4 °C. Membranes were then washed three times with TBS-T followed by incubation during 1 hr at room temperature with anti-rabbit or anti-mouse secondary antibody coupled to HRP (all from Cell signaling) diluted at 1:1,000 in blocking buffer. Membranes were then washed three times in TBS-T. Chemiluminescence was visualized with the Pierce ECL Western Blotting Substrate (Pierce) and acquired on ImageQuant LAS4000 (GE Healthcare). Quantification of band intensity was performed with the ImageJ software. GAPDH or β tubulin was used as loading control and for relative quantification. Positive control for mature IGF2 detection was 500 ng of recombinant mouse IGF2 (R&D Systems).

**Table 4.** List of antibody used for Western Blot.

| Primary antibody | Dilution | Supplier | Clone |
| --- | --- | --- | --- |
| Rabbit anti-IGF2 | 1:500 | AVIVA SYSTEMS BIOLOGY | OAAB07463 |
| Rabbit anti-STAT3 | 1:1000 | Cell Signaling Technology | D3Z2G |
| Rabbit anti-STAT3 (p705 Tyr) | 1:1000 | Cell Signaling Technology | D3A7 |
| Rabbit anti-JNK/SAPK | 1:1000 | Cell Signaling Technology |  |
| Rabbit anti-pJNK | 1:1000 | Cell Signaling Technology | 81E11 |
| Rabbit anti-ERK (1/2) | 1:1000 | Cell Signaling Technology | 137F5 |
| Rabbit anti-pERK (1/2) | 1:1000 | Cell Signaling Technology | Anti-rabbit IgG HRP |
| Rabbit anti-SOCS3 | 1:1000 | Cell Signaling Technology | Anti-rabbit IgG HRP |
| Mouse anti-GAPDH | 1:1000 | Pierce | GA1R |
| Mouse anti-β tubulin | 1:1000 | Pierce | BT7R |
| Mouse anti-VP1 | 1:1000 | Dako | 5-D8/1 |

### Statistical analyses

Statistical analyses were performed with GraphPad Prism 8.0. Unpaired *t* test and ratio paired *t* test were used to compare differences between groups, respectively for *in vivo* and *in vitro* experiments. One-way ANOVA was used to compare differences between time points for *in vivo* and *in vitro* experiments. *P* values inferior or equal at 0.05 were considered significant.

## CONFLICTS OF INTEREST

The authors declare that they have no conflict of interests.

## ACKNOWLEDGEMENTS

This work was supported by doctoral grants obtained from the Fund of Scientific Research (FSR and FRIA of Belgium), by Erasmus+, and by the Fonds Léon Fredericq of Liège University Hospital. The funders had no role in study design, data collection and interpretation, or the decision to submit the work for publication. We thank animal facilities, flow cytometry and also viral vectors platforms from the Institute of Research GIGA at the University of Liège.

